## Supplementary Table 1 and Supplementary Figures for "Exploring the Roles of RNAs in Chromatin Architecture Using Deep Learning"

Supplementary Table 1. Data source used to extract features for AkitaR models.

| Model Input | Experiment | Accession Number | Reference |
| --- | --- | --- | --- |
| Steady State RNA | RNA-seq | 4DNESFH3EHTU | ^45,68^ |
| Nascent RNA | iMARGI | 4DNES9Y1GHK4 | ^20,45^ |
| *Trans*-located RNA | iMARGI | 4DNES9Y1GHK4 | ^20,45^ |
| ATAC-seq | ATAC-seq | 4DNESMBA9T3L | ^45,67^ |

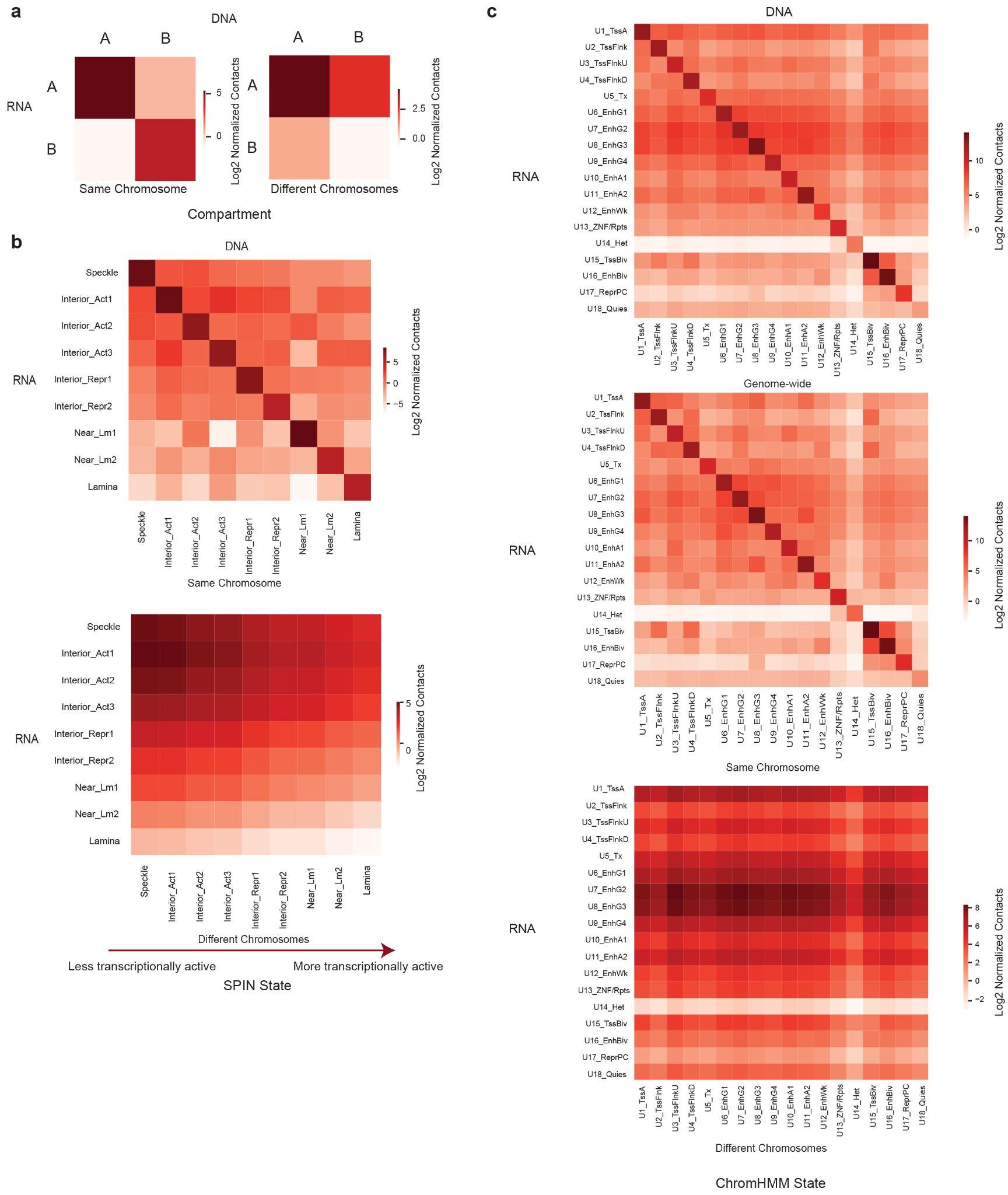

Supplementary Figure 1. RNA-DNA interactions occur across compartments, SPIN states and chromHMM states. The number of RNA-DNA interactions (log_2_) within and across compartments (a), SPIN states (b) and chromHMM states (c). RNA-DNA interactions are separated into the ones on the same chromosome and the ones on different chromosomes. For chromHMM states, genome-wide RNA-DNA interactions combining both the ones on the same and different chromosomes are also shown. Interaction frequencies are normalized to the size of compartments, SPIN states and chromHMM states. U1_TssA: Active TSS, U2_TssFlnk: Flanking TSS, U3_TssFlnkU: Flanking TSS Upstream, U4_TssFlnkD: Flanking TSS Downstream, U5_Tx: Transcription, U6_EnhG1: Genic Enhancer 1, U7_EnhG2: Genic Enhancer 2, U8_EnhG3: Genic Enhancer 3, U9_EnhG4: Genic Enhancer 4, U10_EhnA1: Active Enhancer 1, U11_EnhA2: Active Enhancer 2, U12_EnhWk: Weak Enhancer, U13_ZNF/Rpts: ZNF Genes & Repeats, U14_Het: Heterochromatin, U15_TssBiv: Bivalent/Poised TSS, U16_EnhBiv: Bivalent Enhancer, U17_ReprPC: Repressed PolyComb, U18_Quies: Quiescent

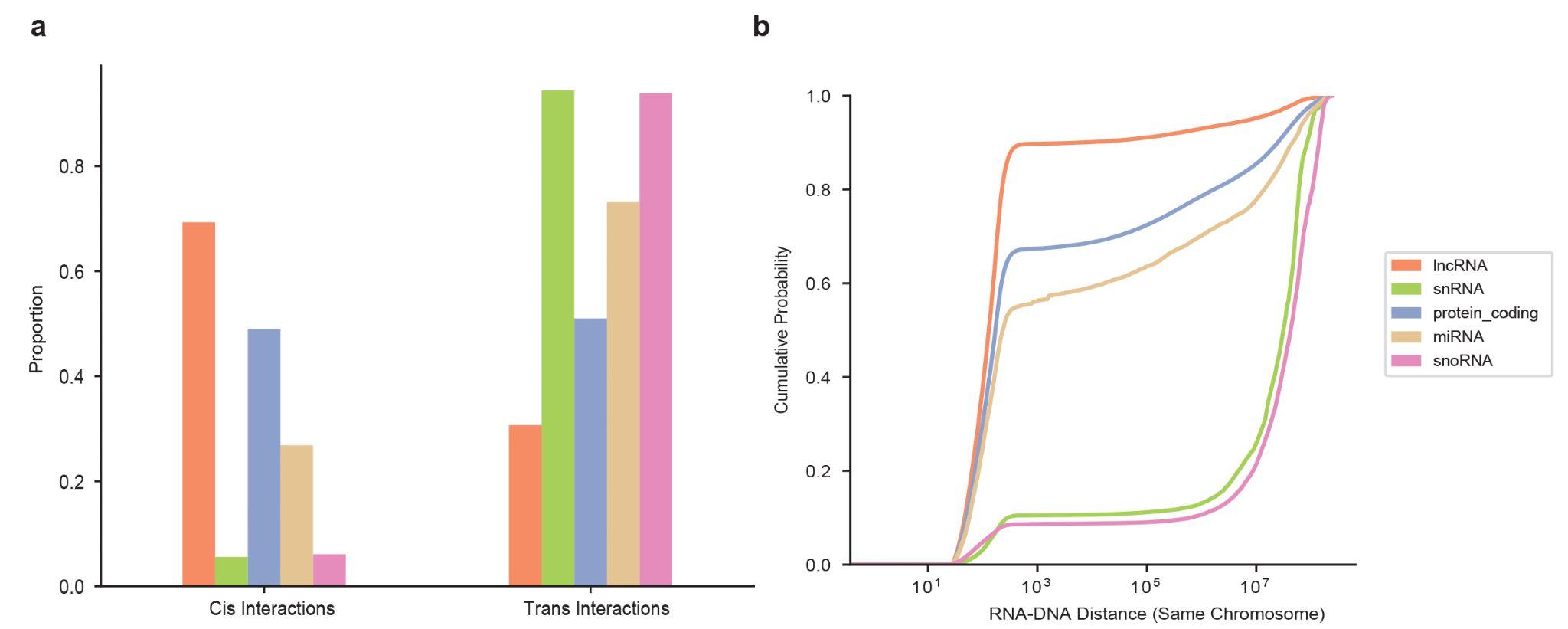

Supplementary Figure 2. RNA-chromatin interactions prevalently occur *in trans*. (a) The proportion of RNAs involved in *trans*-interactions for each RNA type. (b) The cumulative probability of the RNA-DNA interactions as a function of genomic distance between DNA and RNA loci on the same chromosome.

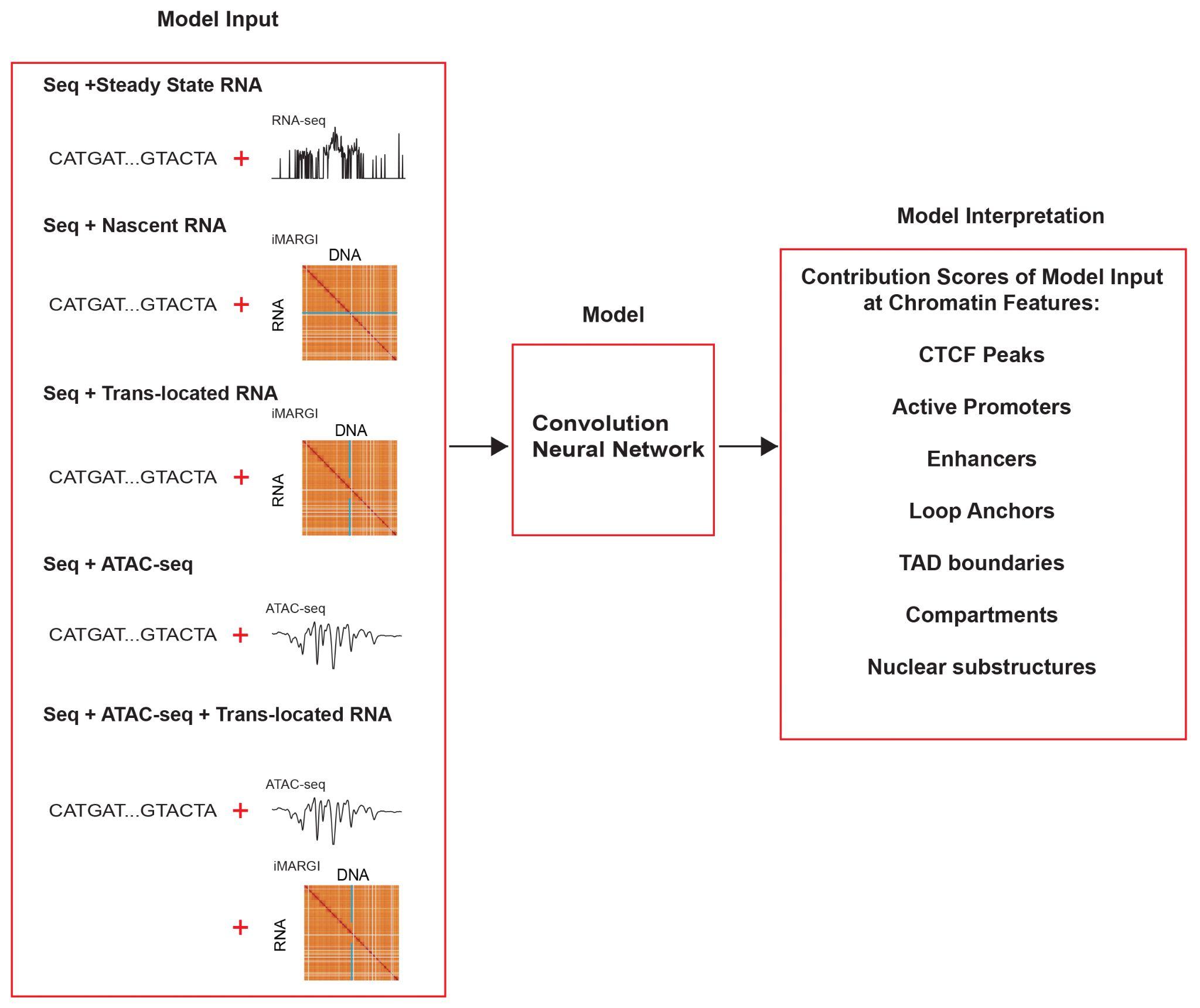

Supplementary Figure 3. Flowchart of workflow for exploring the roles of caRNAs in 3D genome architecture using deep learning. The inputs for the several models trained in this study are shown in the left panel. The chromatin features used to annotate the DNA regions with high contribution scores of different model inputs are shown in the right panel.

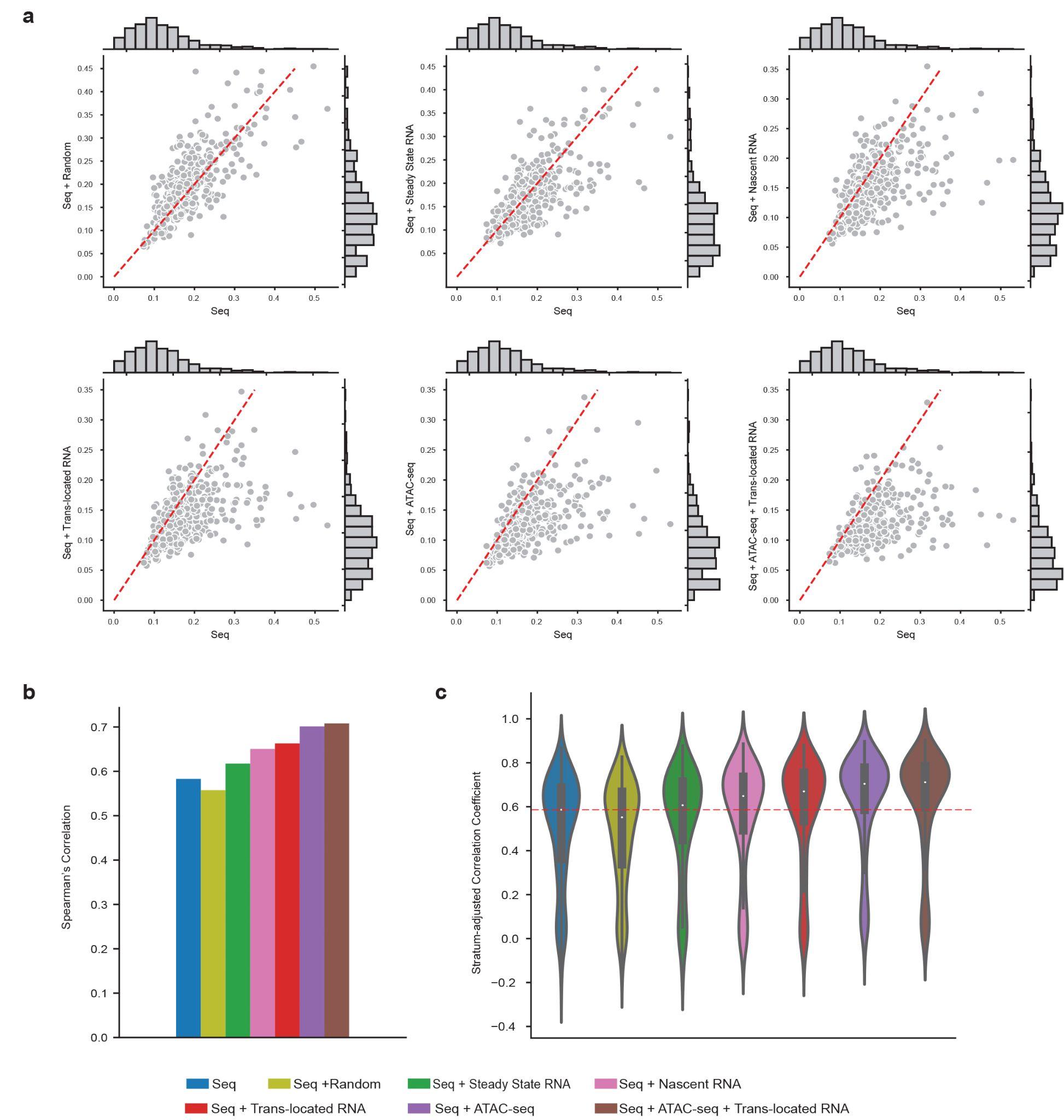

Supplementary Figure 4. Chromatin-associated RNAs increase the accuracy of contact map predictions. (a) Pairwise comparisons of MSE between the models with additional features and sequence alone model. MSE was calculated per held-out test region between experimental and predicted contact maps. The points under the red line (y=x) represent the regions with lower MSE (better performance) for the model with additional features. (b) Barplots of Spearman’s correlation between experimental and predicted contact maps of the held-out test set. (c) Violin plot of stratum-adjusted Pearson’s correlation between experimental and predicted contact maps of the held-out test set.

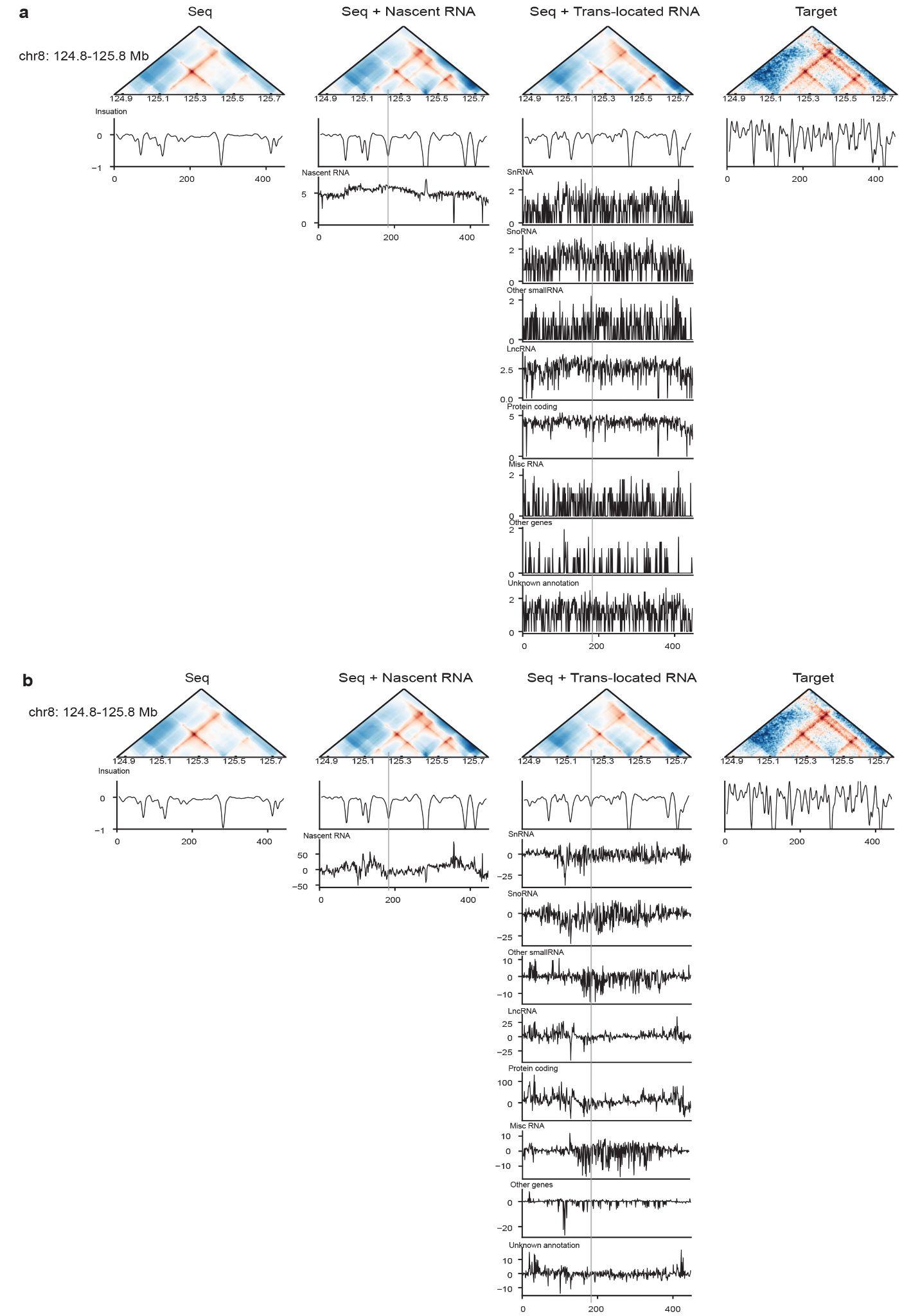

Supplementary Figure 5. An example showing the better predictions of the model with nascent transcription compared to the one with *trans*-located caRNAs. Model inputs and their contribution scores are shown in a and b, respectively. Insulation tracks of the predicted or experimental contact maps were also shown. The region with better predictions by the model with nascent transcription is highlighted with a gray bar.

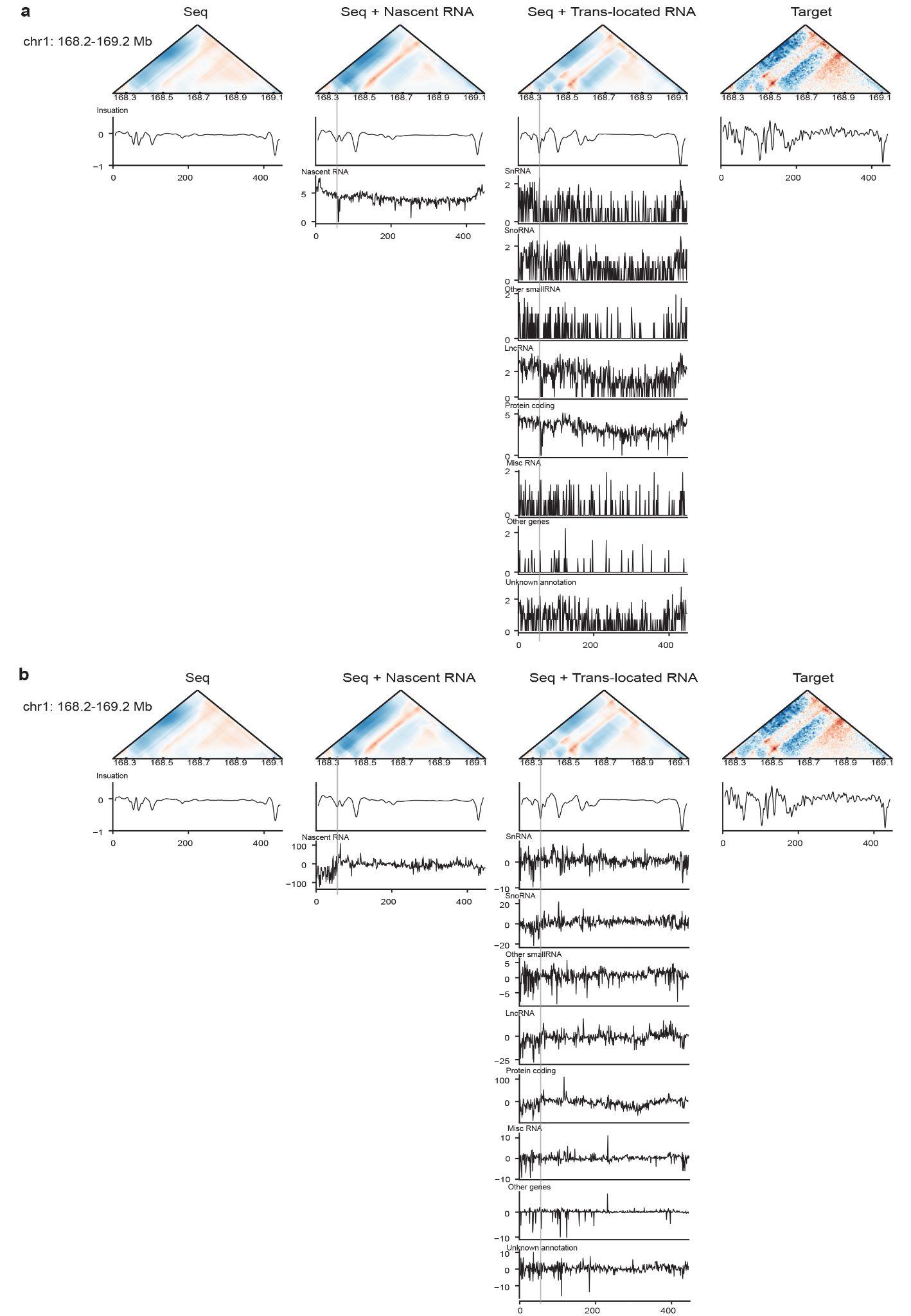

Supplementary Figure 6. An example showing the better predictions of the model with *trans*-located caRNAs compared to the one with nascent transcription. Model inputs and their contribution scores are shown in a and b, respectively. Insulation tracks of the predicted or experimental contact maps are also shown. The region with better predictions by the model with *trans*-located caRNAs is highlighted with a gray bar.

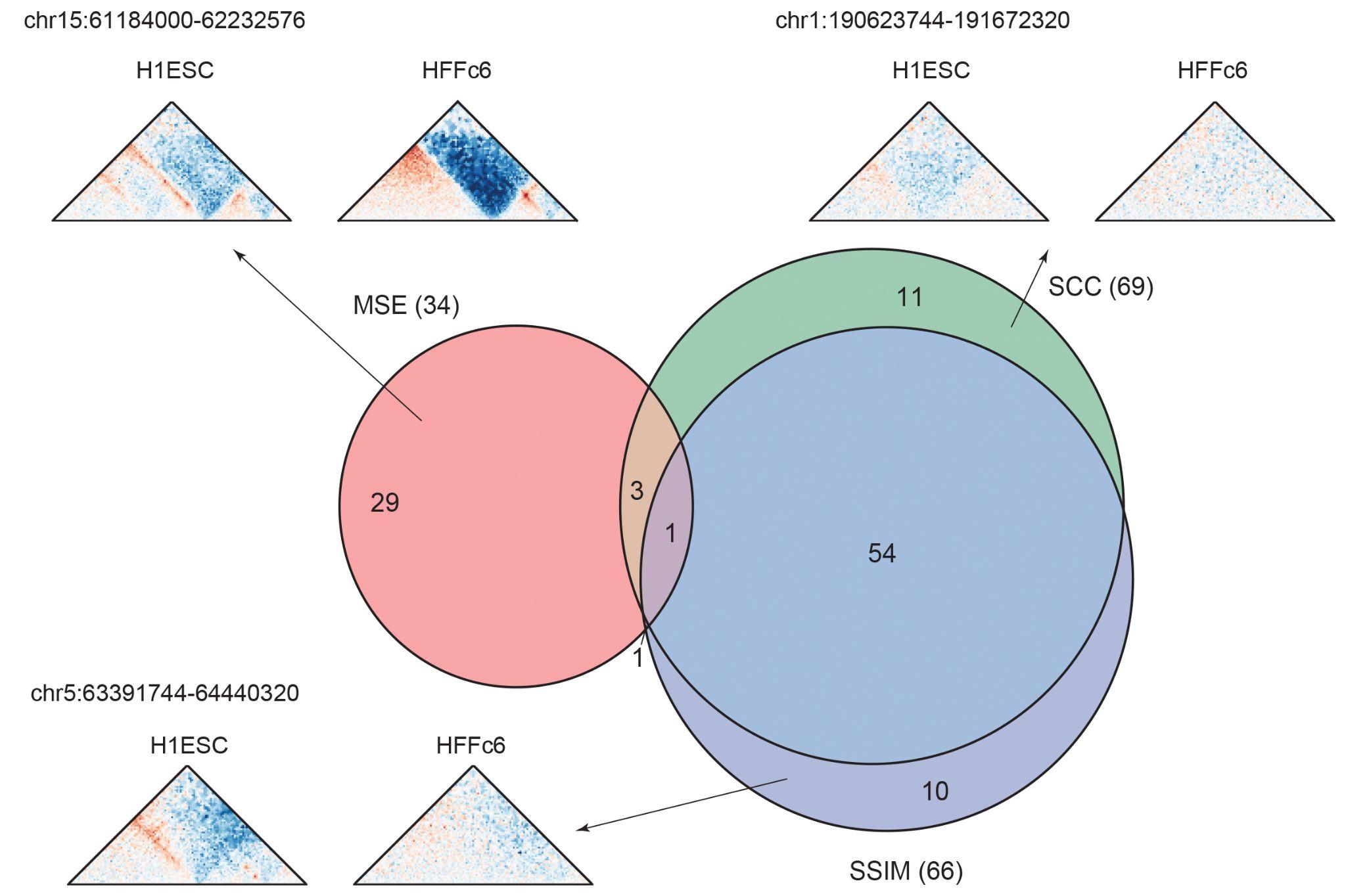

Supplementary Figure 7. Test regions with cell-type differences between Micro-C data in H1ESC versus HFFc6. The cell-type-specific regions were identified by MSE (>0.3), SCC (<0.2) or SSIM (<0.08). A representative example for the cell-type-specific regions identified by each metric is shown.

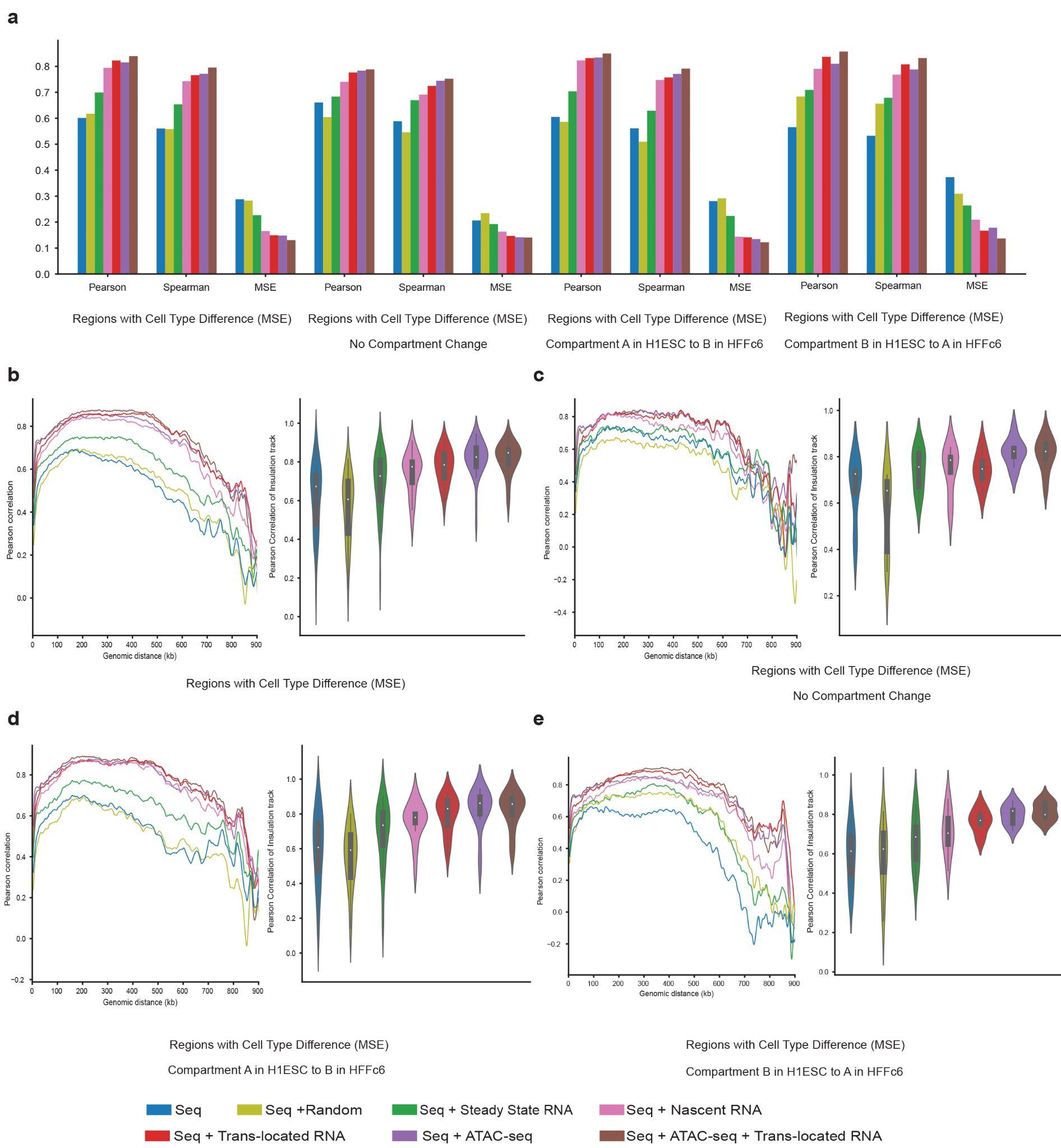

Supplementary Figure 8. Chromatin-associated RNAs help capture cell-type-specific genome folding. (a) Barplots of Pearson’s correlation, Spearman’s correlation and MSE between experimental and predicted contact maps on the cell-type-specific subsets (MSE>0.3 between experimental maps in HFFc6 versus H1ESC) and cell-type-specific subsets (MSE>0.3) without compartment change, with compartment changes from A compartment in H1ESC to B compartment in HFFc6, or with compartment changes from B compartment in H1ESC to A compartment in HFFc6. (b-e) Stratified Pearson’s correlation and violin plot of Pearson's correlation of insulation tracks between experimental and predicted contact maps on the cell-type-specific test subsets identified by MSE (MSE>0.3) (b) and cell-type-specific subsets (MSE>0.3) without compartment change (c), with compartment changes from A compartment in H1ESC to B compartment in HFFc6 (d), or with compartment changes from B compartment in H1ESC to A compartment in HFFc6 (e).

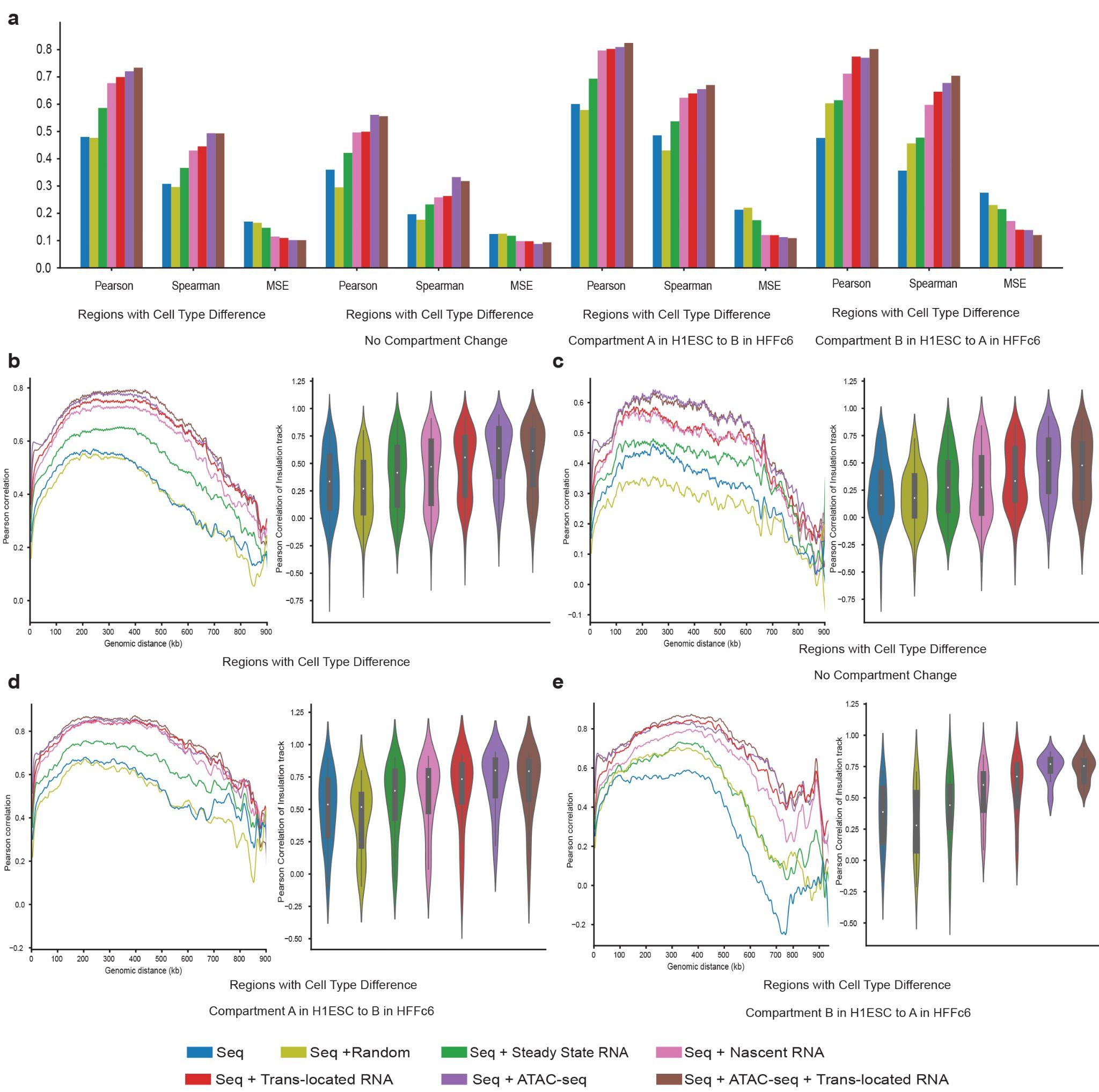

Supplementary Figure 9. Chromatin-associated RNAs help capture cell-type-specific genome folding identified by MSE, SCC or SSIM. (a) Barplots of Pearson’s correlation, Spearman’s correlation and MSE between experimental and predicted contact maps on the cell-type-specific subsets (MSE>0.3, SCC<0.2 or SSIM<0.08) and cell-type-specific subsets (MSE>0.3, SCC<0.2 or SSIM<0.08) without compartment change, with compartment changes from A compartment in H1ESC to B compartment in HFFc6, or with compartment changes from B compartment in H1ESC to A compartment in HFFc6. (b-e) Stratified Pearson’s correlation and violin plot of Pearson's correlation of insulation tracks between experimental and predicted contact maps on the cell-type-specific test subsets (MSE>0.3, SCC<0.2 or SSIM<0.08) (b) and cell-type-specific subsets (MSE>0.3, SCC<0.2 or SSIM<0.08) without compartment change (c), with compartment changes from A compartment in H1ESC to B compartment in HFFc6 (d), or with compartment changes from B compartment in H1ESC to A compartment in HFFc6 (e).

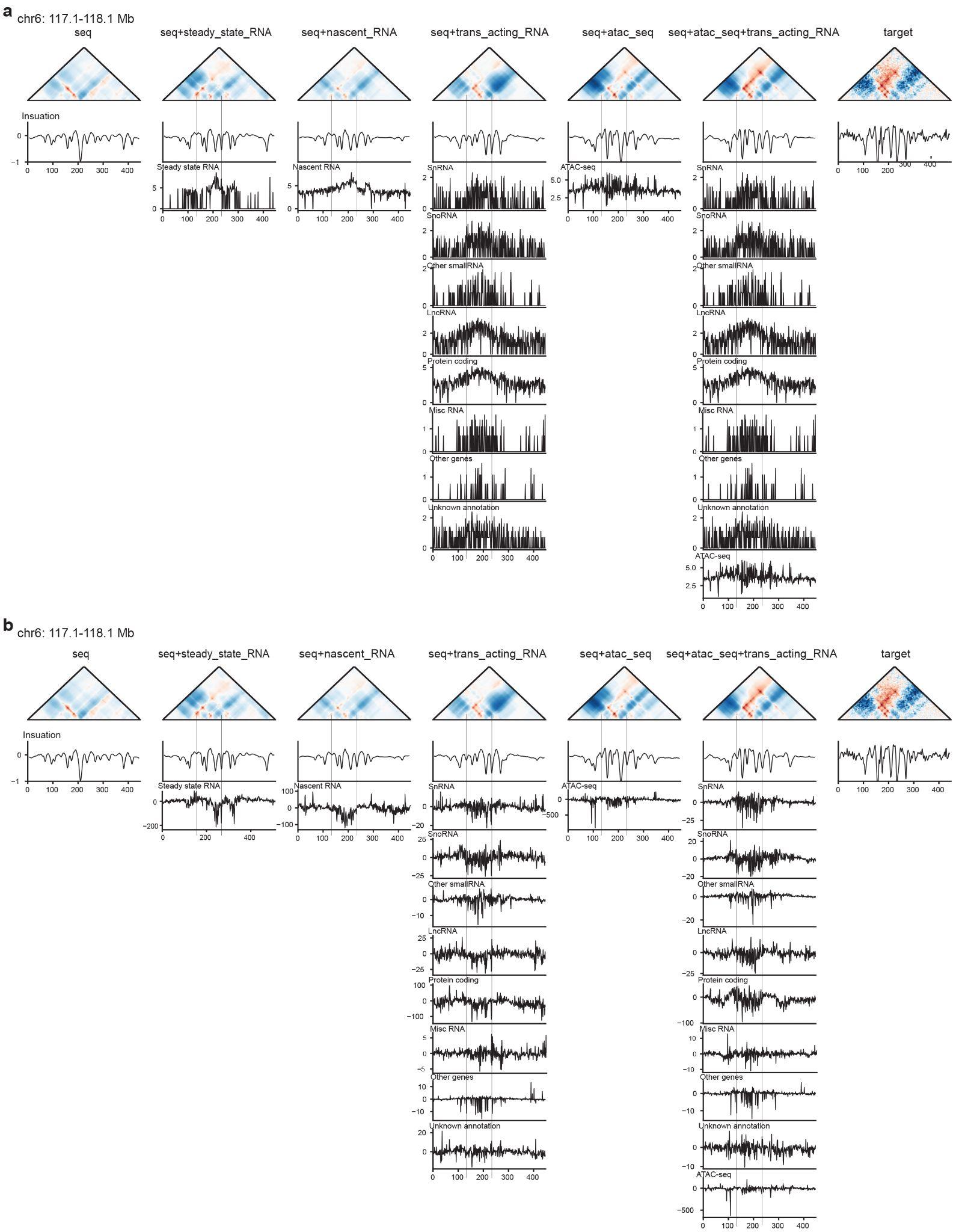

Supplementary Figure 10. An example showing the model with *trans*-located caRNAs captured some cell-type-specific chromatin interactions better than all other RNA and ATAC-seq features. Model inputs and their contribution scores are shown in a and b, respectively. Insulation tracks of the predicted or experimental contact maps are also shown. The region with better predictions by the model with *trans*-located caRNAs is highlighted with a gray bar.

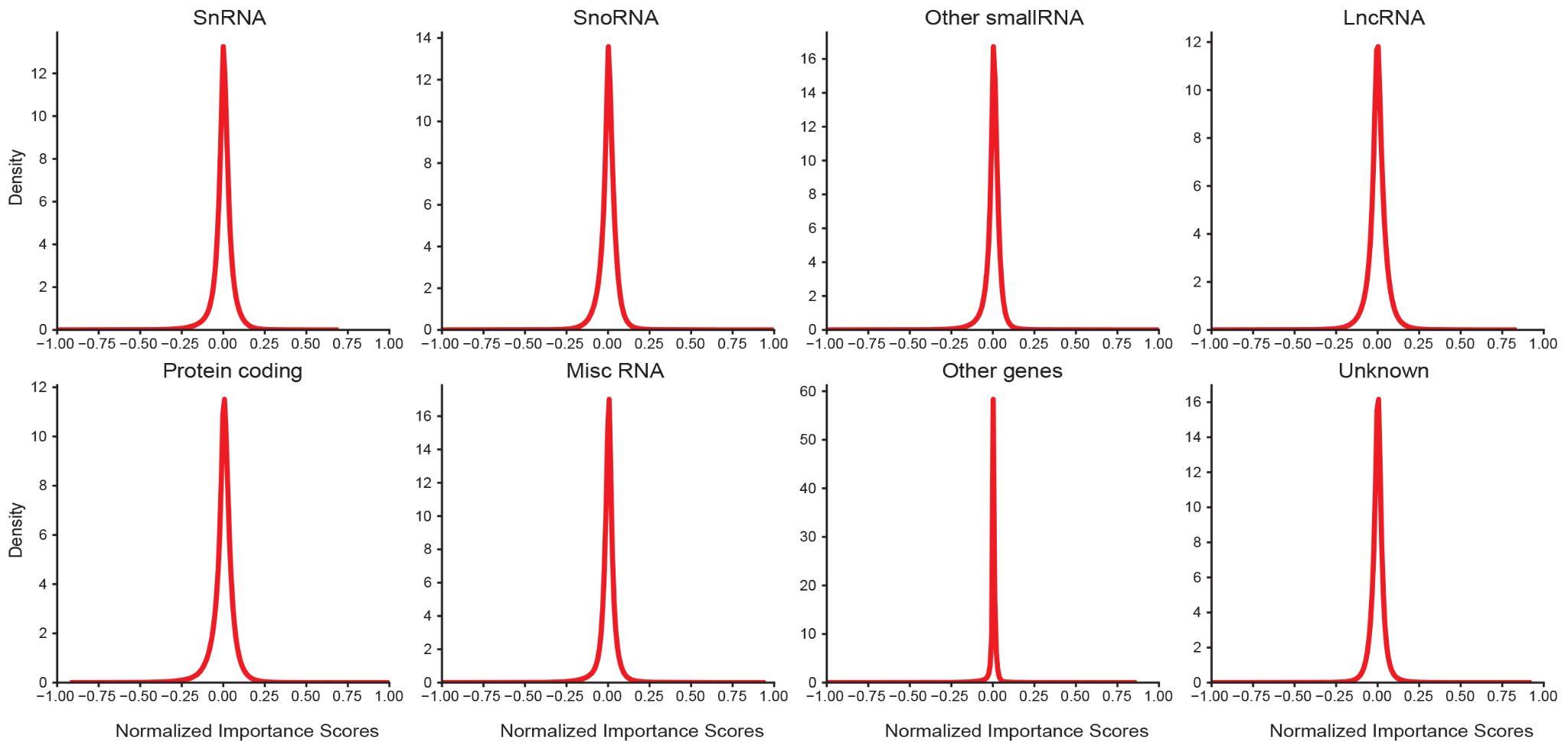

Supplementary Figure 11. Density plot of normalized contribution scores of *trans*-located caRNAs from different RNA types. Contribution scores were normalized to their maximum absolute values. SnRNAs, lncRNAs, RNAs from other types of genes and RNAs from regions without known annotation demonstrated asymmetric distribution with slightly elongated left tails.

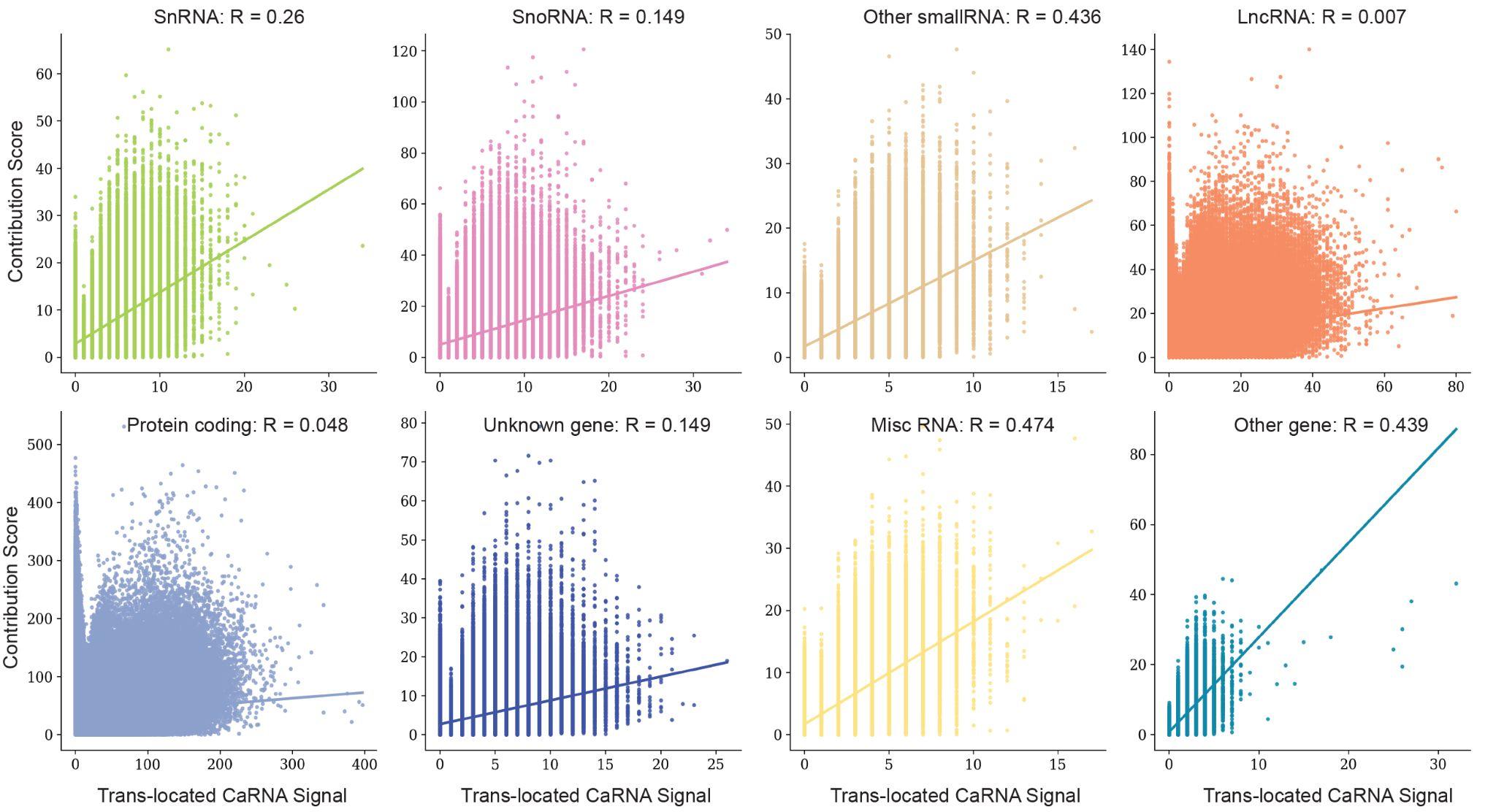

Supplementary Figure 12. Absolute contribution scores of *trans*-located caRNAs from different RNA types show low or medium correlation with their input signals.

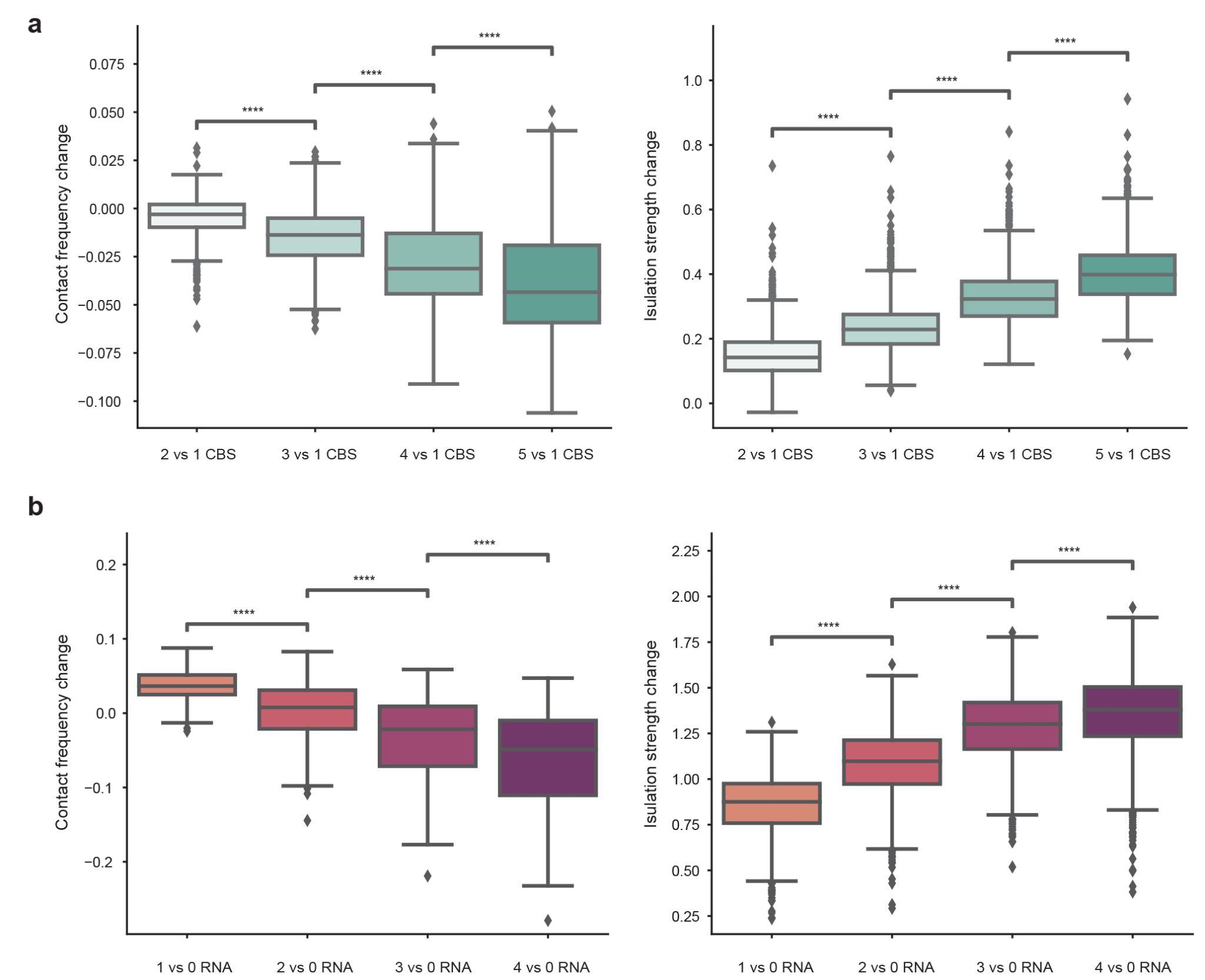

Supplementary Figure 13. *Trans*-located caRNAs might help strengthen the insulation of TAD boundaries. (a) Inserting CTCF motifs progressively from 1 to 4 to random DNA sequence with a pair of convergent CTCF motifs increased the insulation strength and decreased the average contact frequency. Predictions of contact maps were made by the model with sequence alone. CBS: CTCF binding sites. (b) Progressively replacing random caRNAs signals with *trans*-located caRNA signals with large negative scores at TAD boundaries strengthened TAD boundary insulation and decreased the average contact frequency of contact map. Predictions of contact maps were made by the model incorporating DNA sequence and *trans*-located caRNA signals. ****: p-value < 0.0001

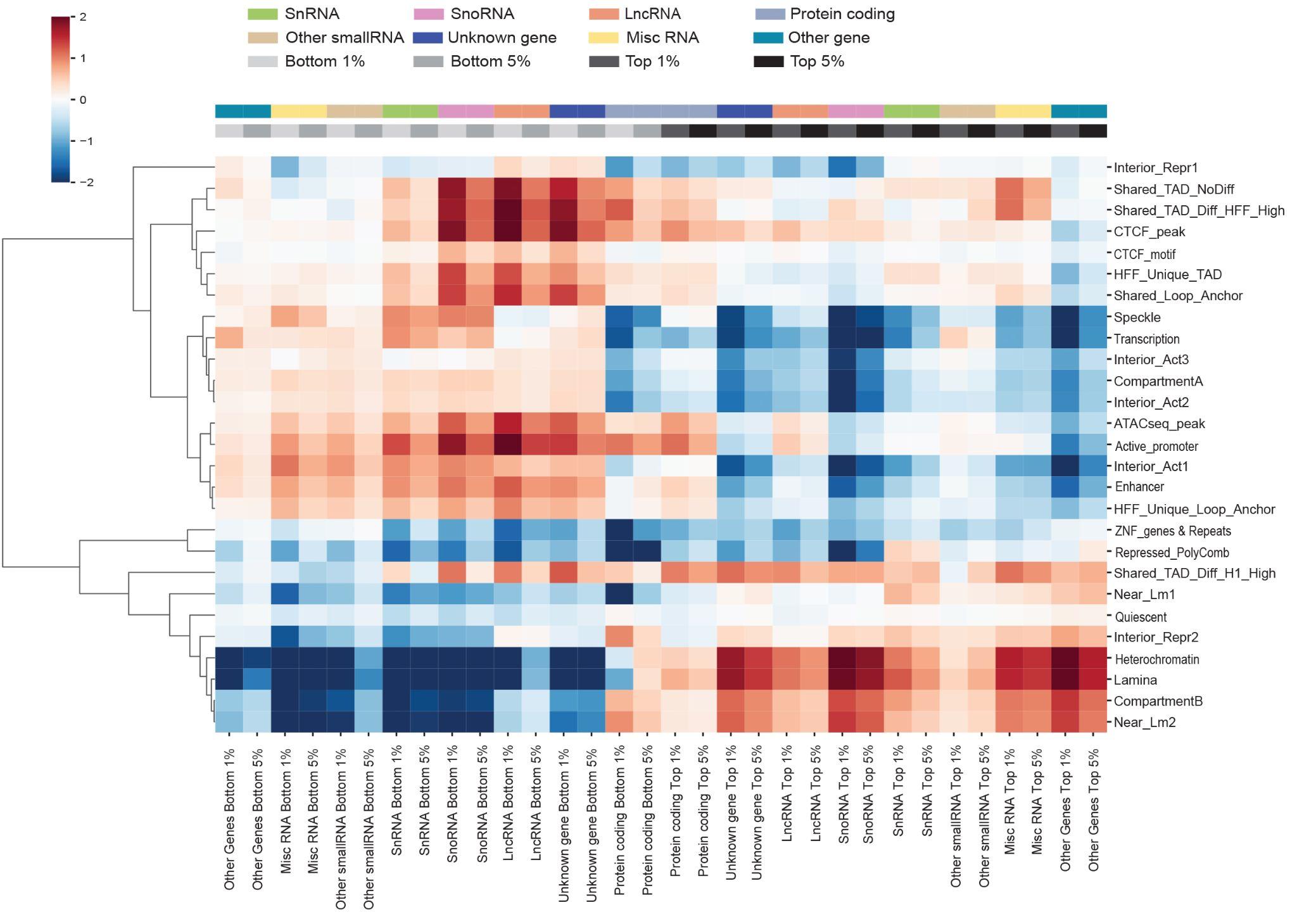

Supplementary Figure 14. Genomic regions with high absolute contribution scores from *trans*-located caRNAs show enrichment at TAD boundaries, loop anchors and nuclear structures. The heatmap shows the enrichment of genomic regions with top 1%, 5% (positive) and bottom 1%, 5% (negative) contribution scores of each type of caRNAs at TAD boundaries, loop anchors, SPIN and ChromHMM states.

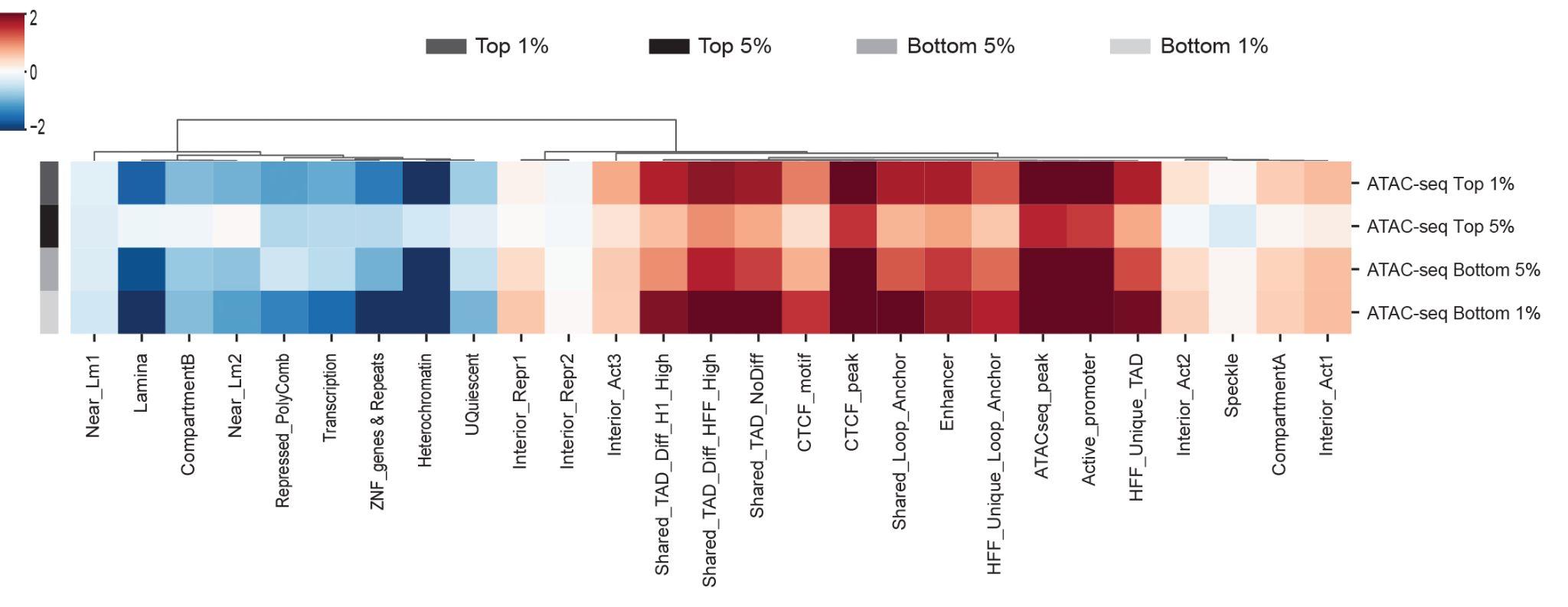

Supplementary Figure 15. Genomic regions with either top or bottom contribution scores from chromatin accessibility show enrichment in active chromatin. The heatmap shows the enrichment of genomic regions with top 1%, 5% (positive) and bottom 1%, 5% (negative) contribution scores of ATAC-seq signals at TAD boundaries, loop anchors, SPIN and ChromHMM states.

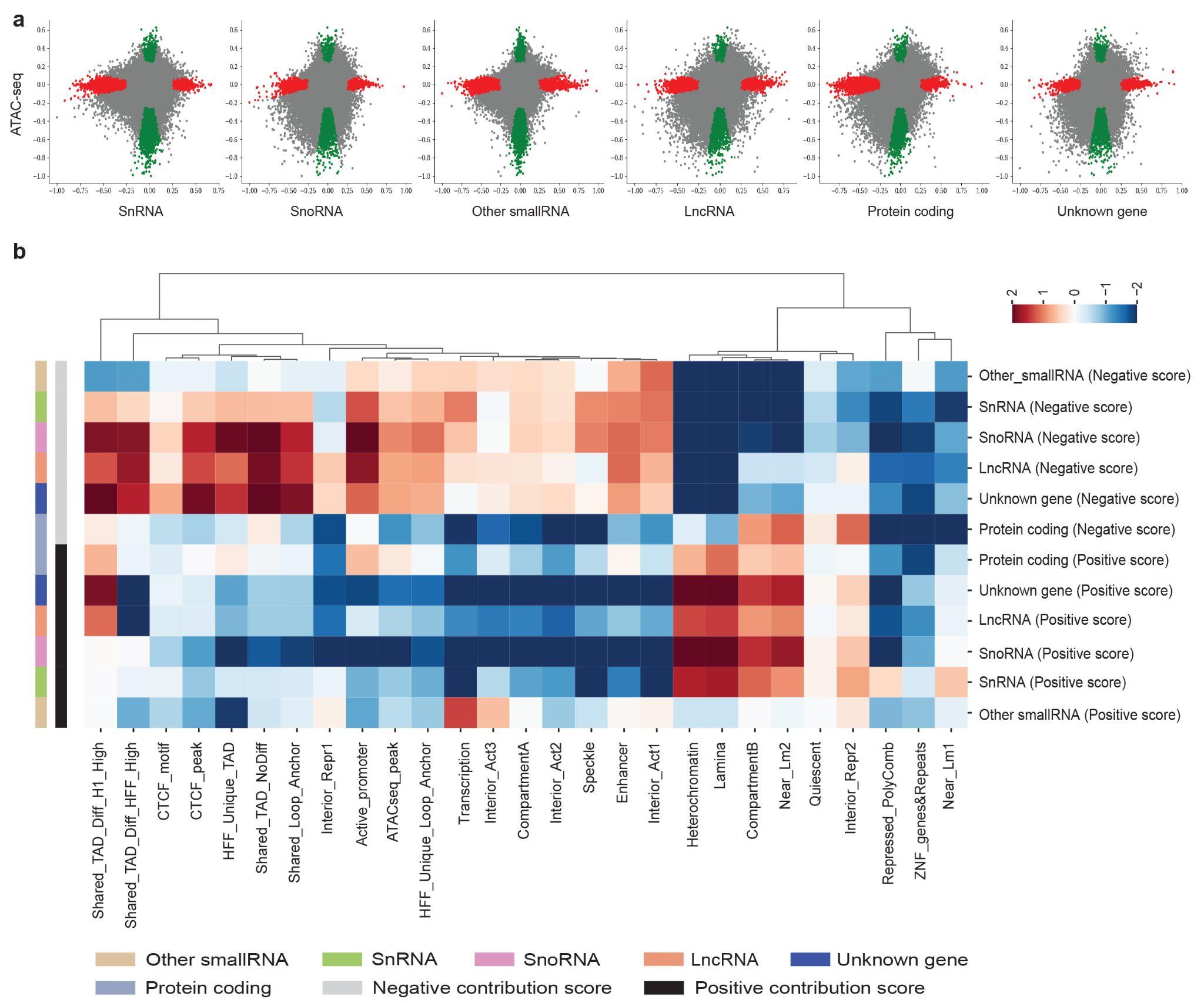

Supplementary Figure 16. Regions with higher contribution scores of *trans*-located caRNAs compared to chromatin accessibility are enriched at TAD boundaries, loop anchors and nuclear structures. (a) Comparison of contribution scores between *trans*-located caRNAs and ATAC-seq signals. (b) Enrichment of genomic regions that showed higher contribution scores for each type of caRNAs compared to ATAC-seq values at TAD boundaries, loop anchors, SPIN and ChromHMM states.

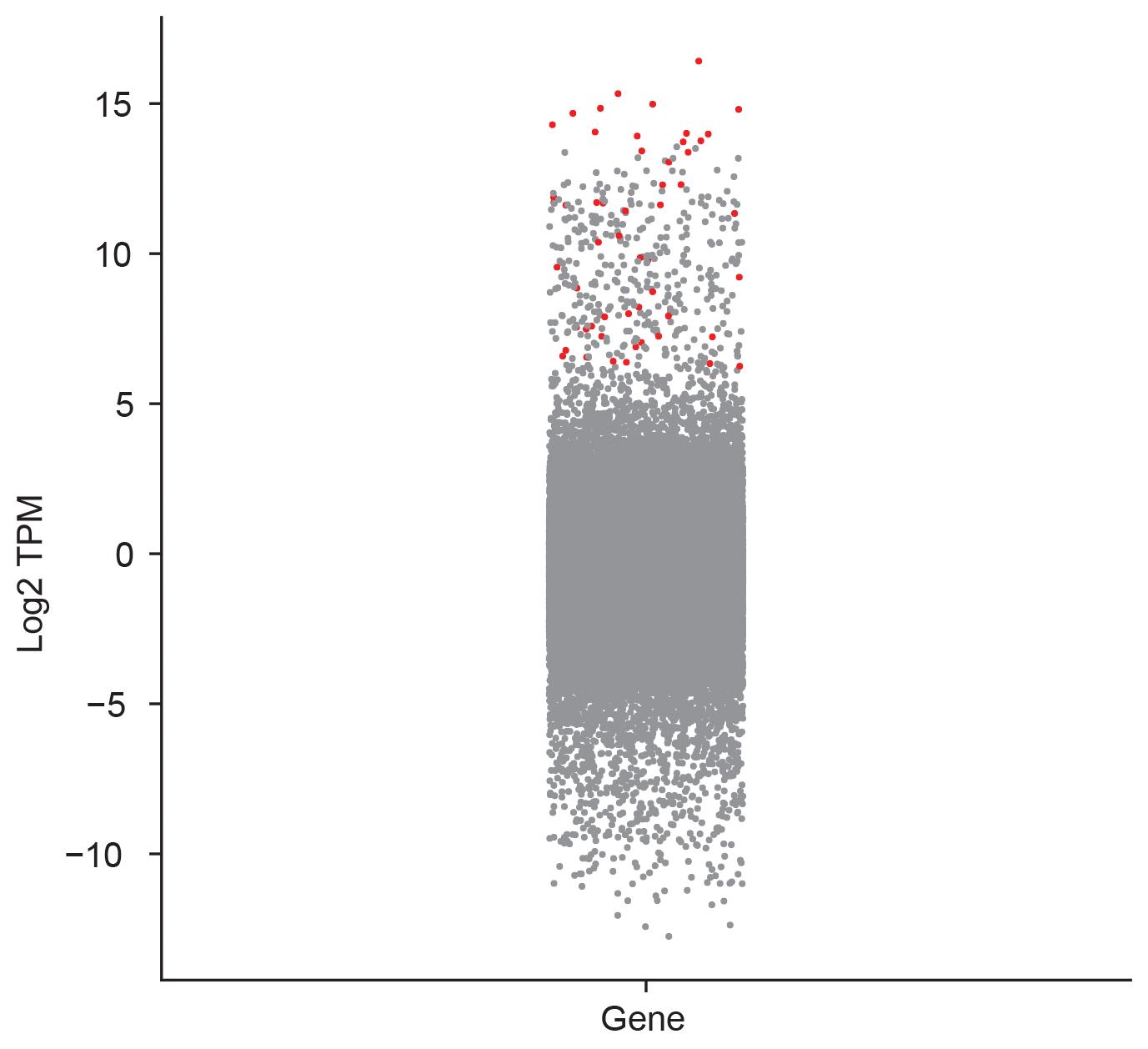

Supplementary Figure 17. Genes whose RNAs preferentially locate at genomic regions with large absolute contribution scores are highly prevalent in HFFc6. Nascent transcription (log_2_TPM) of each annotated gene is shown, and the genes whose RNAs are preferentially located at genomic regions with large absolute contribution scores are highlighted in red.

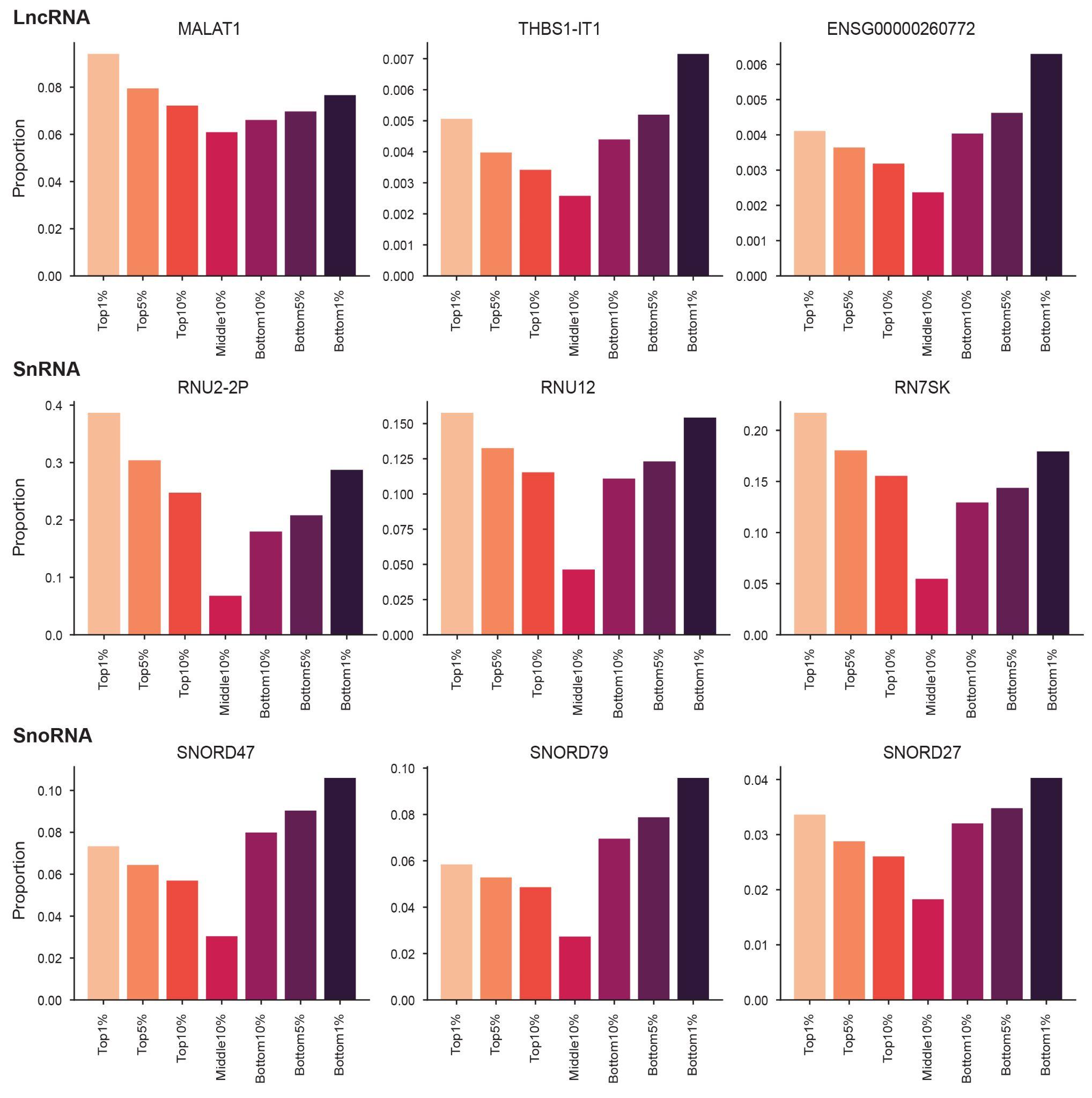

Supplementary Figure 18. Candidate RNAs that are preferentially associated with genomic regions with large absolute contribution scores. Example of lncRNAs, snRNAs and snoRNAs that preferentially interact with genomic regions with large absolute contribution scores (top 1%, top 5%, top 10%, bottom 1%, bottom 5%, bottom 10%) versus regions with lower absolute contribution scores (middle 10%).

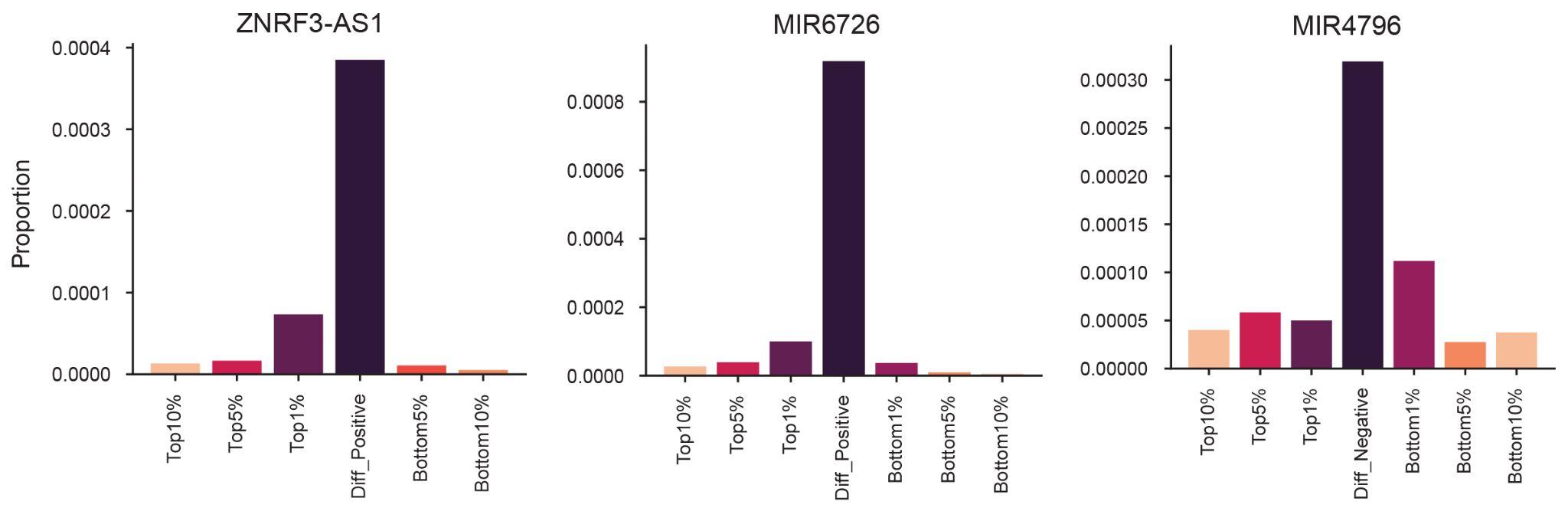

Supplementary Figure 19. Candidate RNAs that are preferentially associated with genomic regions where trans-located caRNAs have large absolute contribution scores and ATAC-seq features do not. Diff_Positive: differentiated regions with larger positive contribution scores for *trans*-located caRNAs. Diff_Negative: differentiated regions with larger negative contribution scores for *trans*-located caRNAs.
